# Build heatmaps of cell frequencies and marker intensities for correct interpretations

**DOI:** 10.1101/2025.02.10.637380

**Authors:** Eugénie Lohmann, Laurent Gorvel, Samuel Granjeaud

**Affiliations:** Centre de Recherche en Cancérologie de Marseille, CRCM, Institut Paoli-Calmettes, Inserm, CNRS, Aix Marseille University, Marseille, France

**Keywords:** cytometry, heatmap, frequency, intensity, method, build, phantasus, heat map, pheatmap

## Abstract

High-content cytometry is an important technique in clinical research, producing data that is rich and complex to analyze. The resulting clusters of cells must be profiled to determine their role and function, and analyzed to identify changes in cell frequency or marker intensity. The heat map is the most suitable tool for presenting this information in a synthetic way. However, its construction must be adapted to the information and the objective. Here we describe the step-by-step construction of heat maps of marker intensity and cell frequency using Excel, Phantasus or R, including the conversion of numbers to colors and the organization of rows and columns. Researchers who master these steps will be able to correctly interpret heat maps and derive the most benefit from high-content cytometry.

## 1 Introduction

The rise of microarray technology in the late 1990s increased the need for visualizing large gene expression datasets to apprehend this data. The article of Eisen et al. (1) established heatmaps as a standard tool for visualization, exploration and analysis. It significantly influenced the field and popularized the use of heatmaps in all omics, as well as the development of software with graphical user interface (2, 3).

In cytometry analyses, heatmaps are used to present in a compact graph the frequency changes of cell clusters (or populations) in the biological samples under study. The aim is to identify frequency changes related to the study question or, less interesting, artefactual effects. Heatmaps are also used to present the expression profile of the cell clusters measured on the markers selected for the study in order to annotate these clusters and associate them with a cell population name.

A heatmap is the representation of a matrix of numbers in an image, using a mapping table that associates a color with a number. A heatmap therefore comprises rows and columns, and the objects they represent must be defined and known. For cytometry, rows and columns may be clusters, markers or samples. The numbers within the matrix are the result of a measurement and its processing, which must be identified in order to make authorized comparisons and avoid misinterpretation. As the color scale is used for all values in the matrix, it is important to know whether comparisons should be restricted within rows or within columns, or not at all. There is no a single magic processing that applies to all types of data. Processing addresses a specific objective and derives from the nature of the measurements and their acquisition. Transforming and scaling are generic terms that compose the processing flow. They require a clear definition and a specific order, which is presented here. Workflows allow to build heatmaps, but their results can be used to apply statistical methods (Principal Components Analysis, etc.) and Machine Learning techniques in order to explore deeply the wealth of cytometry data.

Frequency analysis is based on a matrix of cell counts with rows identifying clusters and columns identifying samples. The aim of the frequency heatmap is to highlight relevant clusters whose frequency changes between groups of samples. The statistical analysis of frequencies is called differential abundance analysis (4) and can be carried out with a dedicated framework such as egdeR (5, 6) or DESeq2 (7).

Cluster profiling is based on a matrix of median fluorescence intensities (MFIs), with rows identifying clusters and columns identifying markers. The term MFI is generic for any flow, mass or spectral cytometry. The aim of the MFI heatmap is to link clusters to known cell populations and complements algorithms such as the MEM method (8).

The MFI analysis of a specified marker is based on a matrix of the MFI values measured per cluster and per sample, with rows identifying clusters and columns identifying samples. In practice, single-marker heatmaps derived from such matrices are rarely presented. However, they can highlight relevant clusters whose MFI changes between groups of samples. More broadly, the aim is to analyze the assemblage of MFI matrices for all markers of interest. This statistical analysis, known as differential state analysis (4), leads to the construction of a meta-heatmap, which integrates data across markers by applying adequate scaling to unify dynamic ranges, and proper normalization to highlight changes in marker intensity.

Being a kind of image, a heatmap should have a carefully chosen color scale to be properly viewed and interpreted. Because cell frequency and marker intensity are on a continuous scale, gradients are the best choice. A single gradient color scale is usually the best option to cover the dynamic range from low to high values. If the mid-point between the minimum and maximum has a specific meaning, a better option is a color scale composed of two diverging gradients, one gradient covering the upper part of the range, and the other the lower part. They meet at the mid-point, where they share the same color. When presenting changes that are symmetric by essence, the mid-point is the “no change” point, and the top and bottom must be symmetric even if the effective data range is not symmetrical. Choosing the colors for the color scale is the final step. There are many options, but it’s important to use common sense (e.g., avoid placing red at the lower end of a blue-white-red scale) and to consider color-blindness.

A heatmap converts numbers into a color matrix, but does not order the rows or columns. When the order is defined by the experimenter or an annotation (such as sample groups), visualization is “supervised”. When the order results from a data-driven algorithm, visualization is “exploratory”. Adding annotations to the rows or columns of the heatmap is useful for visualizing and identifying associations between these annotations and the heatmap data. In the frequency heatmap, samples can be organized into sample groups to facilitate identification of a cluster whose frequency changes are associated to sample groups. But the sample order can also result from a hierarchical clustering, and this natural organization of the data allows us to check whether it correlates with the sample groups. The aim of hierarchical clustering (and other clustering algorithms) is to bring together elements that are similar. Elements are the rows and columns of the heatmap and similarity is based on a distance (also called metric), giving rise to a wide range of options.

There are many approaches to building heatmaps from data organized in a matrix. We use Microsoft Excel as it is widely known and allows each step of the workflows to be easily built and clearly understood. But it does not allow for advanced and necessary final steps such as hierarchical clustering. These final steps are performed using Phantasus (3), a web server with a graphical user interface for transcriptomics data analysis. In addition, we provide the same processing workflows using R and the pheatmap package (9).

## 2 Materials

### 2.1 Software

Requirements are the following:

1. Microsoft Excel, or any spreadsheet software (adaptations are left to the reader).
2. Access to Phantasus website https://artyomovlab.wustl.edu/phantasus/.
3. R and pheatmap package for the alternate R version.

Any recent version of this software will allow you to perform the steps we describe.

### 2.2 Experiment results

The results are taken from a study of Michlmayr et al. (10). They performed a 37-plex mass cytometry acquisition of peripheral blood mononuclear cells (PBMCs) of acute- and convalescent-phase samples obtained from 43 children naturally infected with chikungunya virus. Their analysis classified cells into 57 sub-communities of canonical leukocyte phenotypes and revealed a monocyte-driven response to acute infection, with the greatest expansions in “intermediate” CD14++CD16+ monocytes and an activated subpopulation of CD14+ monocytes. The FCS files and the attachments files are available on FlowRepository under id FR-FCM-Z238. We focused on the smallest subset of cell populations called “Assignment”.

We extracted counts and MFIs from the FCS files although the frequency matrix is available. The extraction and preparation scripts are available in the GitHub repository accompanying this chapter. We used a heatmap of clusters versus samples to show counts lower than 10 cells, which brings a fast overview. We removed clusters with a majority of low counts, reducing the dataset from 30 to 27 clusters. The cell populations differentially abundant in this experiment can be used to verify the heatmaps we obtained and compare them to the original article.

### 2.3 Input data files and format

Cluster profiling is based on a matrix of untransformed MFIs. Rows identify clusters, columns identify markers. The MFI term is generic for any flow, mass or spectral cytometry. Untransformed refers that the MFIs are expressed in the acquisition scale. It is possible to start the workflow with a matrix of transformed MFIs (which corresponds to the second step), and this is a better solution as the transformed MFIs benefit of the transformation tuning performed by the experimenter.

Frequency analysis is based on a matrix of cell counts. Rows identify clusters, columns identify samples (corresponding to FCS files). It is possible to start the workflow with a matrix of frequencies (which corresponds to the second step). We define frequency as the number of cells in a cluster (or a cell population) to the total number of cells in the whole sample (after cleaning steps) or to a selected population (e.g. T cells). Frequency is also called abundancy.

An experiment usually compares samples from different groups. This information as well as preparation and acquisition dates, etc. are important for presenting and interpreting the results and the heatmaps. A metadata file is typically required to link samples to all this information.

We proposed pre-ordered matrices in order to simplify the reading without a hierarchical clustering. Ready to use files are listed in Table 1 and available at https://i-cyto.github.io/build-heatmaps-imp2/.

**Table 1:** Table of available files to reproduce the steps of each workflow.

| **Data file name** | **Description** |
| --- | --- |
| counts_Assignment_matrix.xlsx | Excel sheet to start building the heatmap of frequencies. Also used for DESeq2 analysis. |
| counts_Assignment_matrix-single_sheet.xlsx | Final Excel sheet of processed counts leading to a heatmap. |
| counts_Assignment_matrix-multi_sheet.xlsx | Final Excel sheet of processed counts leading to a heatmap where steps are presented in different sheets. |
| counts_Assignment_matrix-for_phantasus.xlsx | Final Excel sheet of processed counts leading to a log2 relative frequencies matrix for Phantasus. |
| mfis_Assignment_matrix.xlsx | Excel sheet to start building the profiling heatmap. |
| mfis_Assignment_matrix-single_sheet.xlsx | Final Excel sheet of processed MFIs leading to a heatmap. |
| mfis_Assignment_matrix-for_phantasus.xlsx | Final Excel sheet of processed MFIs in a matrix for Phantasus. |
| mfis_samples_Assignment_matrix.xlsx | Excel sheet to start building the meta-heatmap of marker intensity per cluster and per sample. |
| counts_Assignment.RData<br>mfis_Assignment.RData<br>mfis_samples_Assignment.RData | Data files for direct reading in R. |

## 3 Methods

Workflows are summarized in Figure 1, and apply to all cytometry technologies. Cluster profiling is based on the MFIs extracted for each marker of interest. The MFI must be transformed if it is not already. The arcsinh transformation is easy to implement (even in Excel), but the bi-exponential transformation is too complex. Next, the transformed MFIs are normalized to the maximum intensity by marker. Frequency analysis is based on the cell counts measured for each cluster in each sample. The counts are normalized per sample to the total count of the sample, leading to frequencies. The average frequency of each cluster is not interesting in itself, but changes in frequencies within each cluster between samples are important. To capture these changes, the frequencies are normalized per cluster to a reference, usually the average frequency of the cluster. Finally, the logarithm transformation permits to express the increases and decreases in a symmetric way, but to avoid issues with zero frequencies (resulting from zero counts) a threshold can be applied beforehand. Other solutions exist to overcome the problem of the logarithmic transformation of zero (see Note 1).

**Figure 1:**
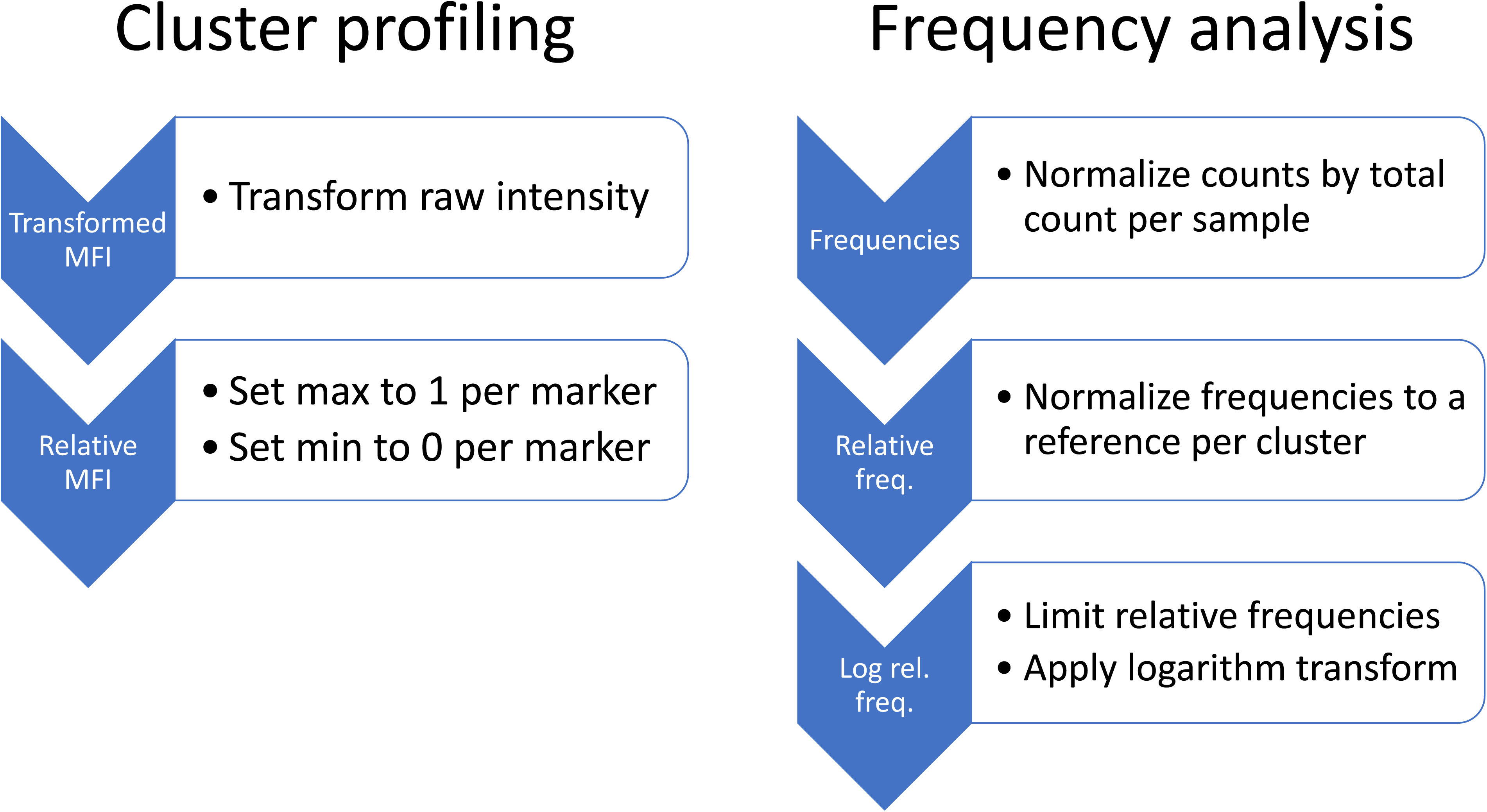
Summary of the main steps of each workflow.

Phantasus can carry out the complete workflow from raw MFIs to heatmap but with some cautions (see Note 2). It allows to perform statistical analyses (see Note 3). It can carry out the complete workflow from counts to heatmap but with some cautions when some counts are zero (see Note 4). Adding a small value to counts is presented with Phantasus (for frequencies, see Note 5).

Here we report protocols built under Windows and an English version of Excel. See Note 6 for Apple and Excel internationalization and Note 7 why Excel file format is interesting.

### 3.1 Building the heatmap of cluster MFI using Excel and Phantasus

#### 3.1.1 Preparing

Make a copy of the MFI matrix Excel file and give it a name. Then open it. Enlarge the clusters column. Reduce the width of the marker columns to fit them all into the screen.

#### 3.1.2 Transforming MFI

The MFI are not transformed, which could be identified by looking at the minimum and maximum of all MFIs. Transformed MFIs are usually below 10. Select and copy the entire region (Ctrl-* Ctrl-C). Go to A31, 3 lines below the copied region and paste (Ctrl-V). This region will hold the transformed MFIs. Go to B32, the top left MFI, and enter the formula “`=ASINH(B2/5)`”. This will transform all the MFIs with the asinh transformation and a co-factor of 5, which is adapted for mass cytometry. Copy the formula to right up to the last marker and down to the last cluster.

#### 3.1.3 Scaling MFI

We will compute the minimum and maximum of each marker (column). Go to A60 and enter “Maximum”. Go to B60, enter the formula “`=MAX(B32:B58)`” and copy to the right up to the last marker. Go to A61 and enter “Minimum”. Go to B61, enter the formula “`=MIN(B32:B58)`” and copy to the right.

Go to A31, select and copy the entire region (Ctrl-* Ctrl-C). Go to A64, 3 lines below the min and max rows and paste. This region will hold the scaled MFIs. Go to B65, the top left MFI, and enter the formula “`=(B32-B$61)/(B$60-B$61)`”. This will scale all the transformed MFIs per marker. Copy the formula to right up to the last marker and down to the last cluster.

#### 3.1.4 Coloring and adjusting

Go to B65 and select the scaled MFIs region. In the “Conditional Formatting” menu, in the “Color Scales” sub-menu, select the “Red-White-Blue” color scale (red being the highest).

To hide the numbers in the cells, select the colored region, then click on the right button of the mouse to format the cells. In the cell format, select “Custom” and in the “Type” of the format, enter “;;;”.

The final heatmap is presented on Figure 2A. As the clusters are already annotated and organized in major cell populations, the heatmap permits to verify that the cluster annotations match the color profiles and that the workflow creates an informative and useful visualization.

**Figure 2:**
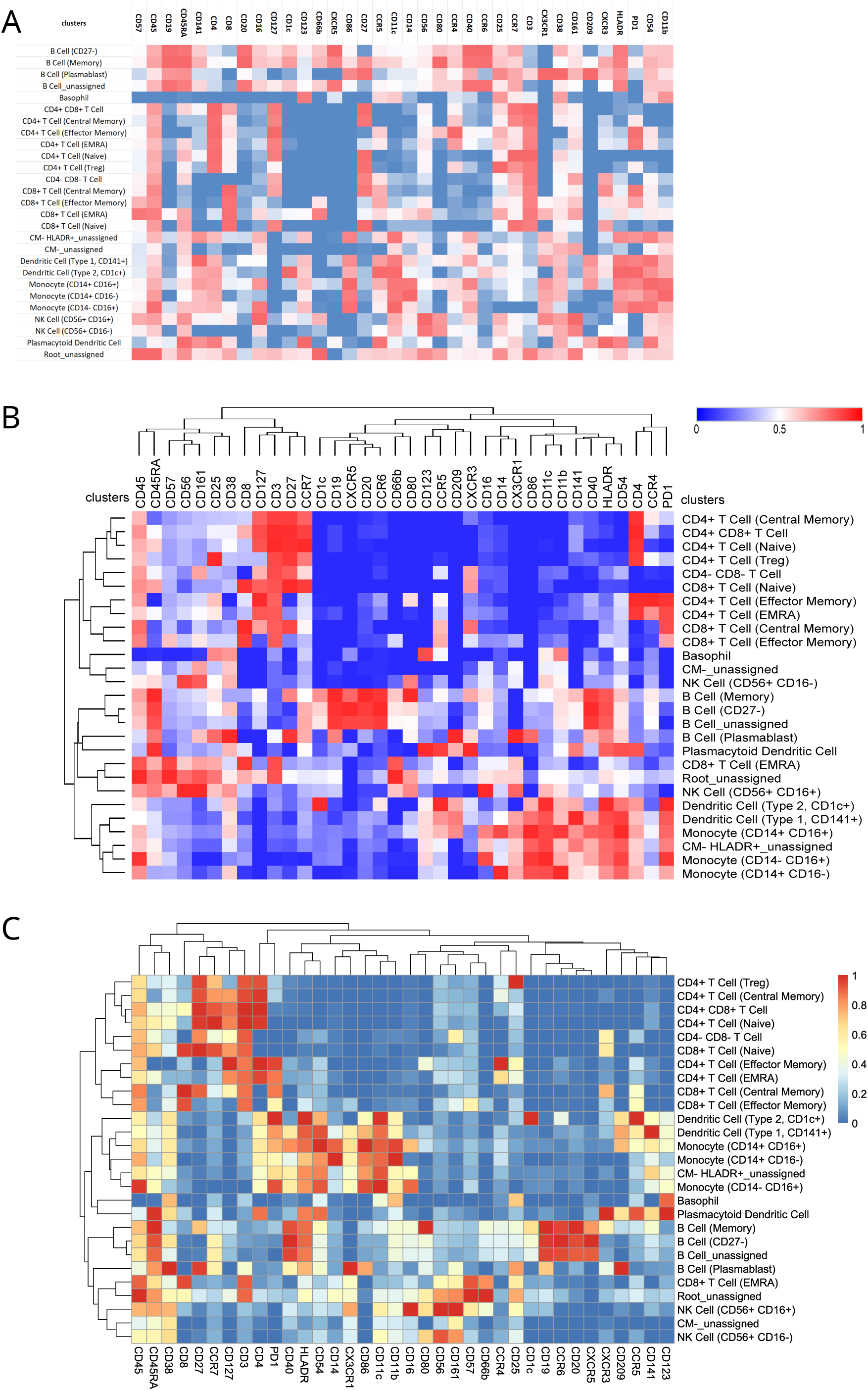
Final heatmaps of transformed MFIs to profile the clusters. A) Heatmap from Excel. B) Heatmap from Excel and Phantasus with clustering of rows and columns to ease the interpretation by patterns. C) Heatmap from pheatmap with clustering.

We can modify the minima and maxima to adjust the coloring to the expected color. For example, the minimum of CD45 is nearly as high as the maximum because all cells were selected to be CD45+. However, the current coloring suggests that low CD45 clusters exist. The minimum intensity should be adjusted to zero, as zero is the expected minimum for CD45. This calibration ensures all clusters are rendered CD45+. A similar adjustment should be performed with maxima of lowly expressed markers such as PD1 that appears too high in many clusters.

#### 3.1.5 Exporting from Excel to Phantasus

Phantasus can import Excel files. See in Note 7 for reasons why using XLSX (Excel files) is a better format than CSV (Comma Separated Values files). We are going to copy the scaled MFI region to a new Excel file. As the colored heatmap region does not show values, Phantasus cannot find values and we have to set up a region with the scaled values.

Go to A64, select the region of the heatmap (Ctrl-* Ctrl-C). Go to A94, 3 lines below the heatmap region, and paste. This region will hold the scaled MFI values. Go to B95, the first MFI value, and enter the formula “=B65”. Copy it to the right and to the bottom of the region.

This formula copies the scaled values. Remove the customized format and set it to “Number”. Values are now visible inside the colored cells. Select the region (Ctrl-*), copy it (Ctrl-V), start a new Excel file and paste only values (an example file is available).

Go to Phantasus website, then select “My computer” in the “Choose a file” pop menu. If Phantasus is already opened in your web browser, click on “File/Open” to import a file. Select the prepared Excel file in your computer. Phantasus presents the annotations and the numerical data matrix of the file. Rows annotations are highlighted in light green and are on the left of the sheet. Column annotations are highlighted in light red and are at the top of the sheet. The data are highlighted in light blue. Click the first cell containing data if the MFIs region is not correctly identified. It is possible to transpose the matrix (i.e. swap rows and columns). Here we consider clusters are on the rows and markers are on the columns. Click “OK”.

To tune the display to the effective scale of the data, open the wheel called “Options” on the top-right. Untick “Relative color scheme” that is using the minimum and maximum on each row to adjust the color scale per row. Verify that “Transform values” is set to “None” (more about on-the-fly transformations in Note 8), “Minimum” to “0” and “Maximum” to “1” (or to “-0.1” and “1.1” to lower the contrast of the color scale). This setting leads to minor changes in the display, but is the only way to show correctly the scaled MFIs. Before closing the window, it is possible to store this color scheme for reuse. Use the “Save color scheme” at the button to specify a name for this color scheme. It will be available in the “Saved color scheme” popup menu; select it and click “Load color scheme” to apply it.

#### 3.1.6 Clustering with Phantasus

Clustering the scaled MFIs really eases the reading and interpretation of the heatmap in order to annotate. The ordering of clusters and markers will only be based on the scaled MFI values.

To apply a clustering, click the “Tools” menu, then move the cursor down to “Clustering”, then select “Hierarchical clustering” in the sub-menu. In the popup window, select “Euclidean” as “Metric” and “Complete” as “Linkage method” (this linkage make clusters more homogeneous). Select the combination of rows and/or columns to which these settings will be applied to. Click “Close” to apply. The final heatmap is presented on Figure 2B (see Note 9 for exporting).

### 3.2 Building the heatmap of cluster frequency using Excel and Phantasus

#### 3.2.1 Preparing

Make a copy of the count matrix Excel file and give it a name. Then open it. Format the cluster names column, the sample names row and the width of count columns.

#### 3.2.2 Computing frequencies

Go to A30, 2 lines below the bottom of the count region. This line will hold the total counts. In A30, enter “`Total`”. In B30, enter the formula “=SUM(B2:B28)” which computes the sum of counts for the first sample. Drag the formula to the right up to the last sample to copy it.

Go to A01, select all counts including sample names and cluster names by typing keys Ctrl-*, then copy the region with Ctrl-C.

Go to A33, 5 lines after the end the counts region and paste by pressing Ctrl-V. This region will hold the frequency of clusters.

Go to B34, the top-left value of the pasted region. In B34, enter the formula “=B2/B$30” which divides each count by the total count of its sample. Copy the formula to the right up to the last sample and to the bottom up to the last cluster.

#### 3.2.3 Computing relative frequencies

Go to CK33, 2 columns after the last sample of the frequency region. A new column will hold the average frequency of each cluster. Enter “Ave frequency”. Go to the cell below at CK34.

`Enter the formula “=AVERAGE(B34:CI34)”`. Copy the formula down to the last cluster.

Go to A33, select all counts including sample names and cluster names and copy the region using Ctrl-*, then Ctrl-C.

Go to A63, 3 lines below the frequencies region and paste. This region will hold the relative frequencies. Go to B64. Enter the formula “=B34/$CK34” which divides the frequency by the average frequency of its cluster. Copy the formula to the right up to the last sample and to the bottom up to the last cluster.

#### 3.2.4 Computing logarithm of limited relative frequencies

Go to A93, 3 lines below the relative frequency region. Enter “Threshold”. Go to B93, enter its value, i.e. the number “4”. This value can be tuned to adjust the visual rendering.

Go to A63, select all relative frequencies including sample names and cluster names and copy the region (Ctrl-* Ctrl-C).

Go to A95, 2 lines below the threshold and paste. The new region will hold the logarithm of threshold relative frequencies. In B95, enter the formula “=LOG(IF(B64>$B$93; $B$93; IF(B64<1/$B$93; 1/$B$93; B64)))/LOG(2)”. The external part of the formula computes the logarithm in base 2. The internal part of the first logarithm function aims to limit values higher than threshold or lower than 1/threshold. Copy the formula to the right up to the last sample and to the bottom up to the last cluster.

#### 3.2.5 Coloring

Go to A95, select all counts including sample names and cluster names and copy the region (Ctrl-* Ctrl-C).

Go to A125, 3 lines below the ready to use values. The new region will hold the colorized version of these values. First paste the copied region, then paste the values only. This will keep the formatting and the values but not the formula.

In the “Conditional Formatting” menu, in the “ Color Scales” sub-menu, select the “Red-White-Blue” color scale (red being the highest).

To hide the numbers in the cells, select the colored region, then click on the right button of the mouse to format the cells. In the cell format, select “Custom” and in the “Type” of the format, enter “;;;”. The final heatmap is presented on Figure 3A.

**Figure 3:**
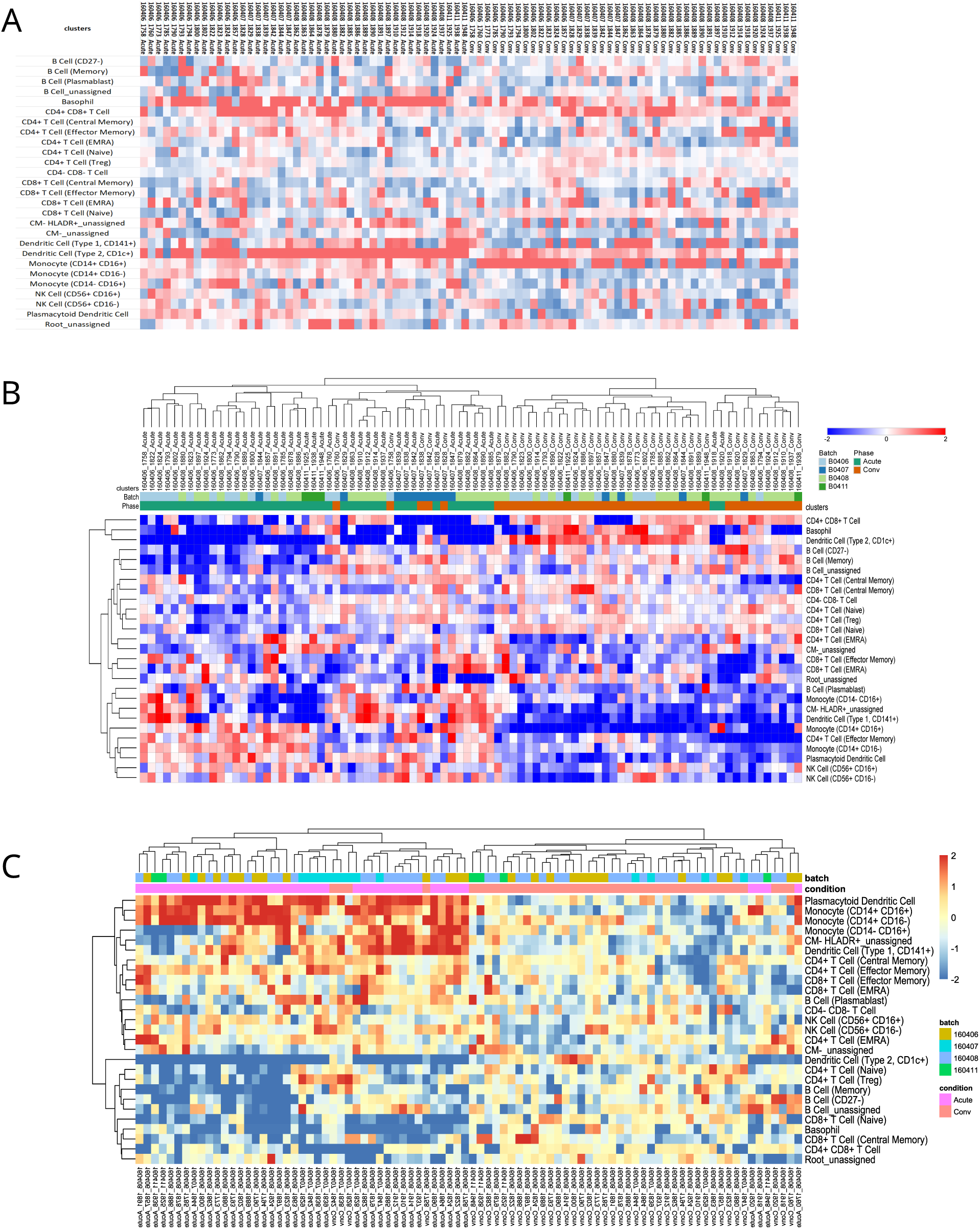
Final heatmaps of relative frequencies in log scale. A) Heatmap from Excel. B) Heatmap from Excel and Phantasus with sample annotations and clustering of rows and columns. C) Heatmap from pheatmap with sample annotations and clustering of rows and columns. Frequencies are relative to the “Conv” sample, making it easier to interpret changes and their significance, and to assess the residual dispersion of the “Conv” sample group.

#### 3.2.6 Exporting from Excel to Phantasus

Phantasus can import Excel files. See in Note 7 for reasons why using XLSX (Excel files) is a better format than CSV (Comma Separated Values files). Start a new Excel file and copy the log2 limited relative frequencies region with values (an example file is available). The colored heatmap region is not recognized as data by Phantasus. Annotations to samples could be added as rows below the first row that contains the sample names. In the available example, we added rows for the group and date given in FCS file names.

Open the website, then select “My computer” in the “Choose a file” pop menu. Select the prepared Excel file in your computer. Phantasus presents the annotations and the numerical data matrix of the file. Rows annotations are highlighted in light green and are on the left of the sheet. Column annotations are highlighted in light red and are at the top of the sheet. The data are highlighted in light blue. Click the first cell containing data if there are annotations. It is possible to transpose the matrix (i.e. swap rows and columns). Here we consider clusters are on the rows and samples are on the columns. Click “OK”.

To tune the display to the effective scale of the data, open the wheel called “Options” on the top-right. Untick “Relative color scheme” that is using the minimum and maximum on each row to adjust the color scale per row. Verify that “Transform values” is set to “None” (more about on-the-fly transformations in Note 8). Set the “Minimum” to “-2” and the “Maximum” to “+2”. Before closing the window, it is possible to store this color scheme for reuse. Use the “Save color scheme” at the button to specify a name for this color scheme. It will be available in the “Saved color scheme” popup menu; select it and click “Load color scheme” to apply it.

#### 3.2.7 Clustering with Phantasus

To apply a clustering, click the “Tools” menu, then move the cursor down to “Clustering”, then select “Hierarchical clustering” in the sub-menu. In the popup window, select “Euclidean” as “Metric” and “Complete” as “Linkage method” (this linkage make clusters more homogeneous). Select the combination of rows and/or columns to which these settings will be applied to. Click “Close” to apply. The final heatmap is presented on Figure 3B (see Note 9 for exporting).

### 3.3 Building the heatmaps with R and pheatmap package

We provide RMarkdown scripts and their corresponding HTML results to present the steps in R programming language. The codes are accessible with a little knowledge of R language.

The aim is to clarify the steps and permit to build a reusable and reproducible workflow to create heatmaps. The pheatmap (9) function is particularly pleasant to create heatmaps and does not carry out on-the-fly transformation by default. It permits to define the limits of the color scale, which avoids the need of thresholding data. It also performs a hierarchical clustering by default without taking into account of the limits applied for the color scale. In this section we reduce the codes to the core formulas (see Note 10 for loading prepared data files).

#### 3.3.1 Heatmap of scaled MFIs

Once the MFIs matrix is loaded (clusters on rows, markers on columns), the workflow is performed with the following code:

mfis = asinh(untransformed_mfis / 5) # cofactor for mass cytometry mfis_max = apply(mfis, 2, max) # max per column, 2nd dimension mfis_to_max = sweep(mfis, 2, mfis_max, “/”) # normalize per column pheatmap(mfis_to_max)

To define a minimum per marker and to limit the color scale between 0 and 1, the code is below:

`mfis_min = apply(mfis, 2, min)`

`mfis_to_min_max = sweep(mfis, 2, mfis_min, “-”) mfis_max = apply(mfis_to_min_max, 2, max)`

`mfis_to_min_max = sweep(mfis_to_min_max, 2, mfis_max, “/”) pheatmap(mfis_to_min_max, breaks = seq(0, 1, length.out = 101))` The final heatmap is presented on Figure 2C.

#### 3.3.2 Heatmap of relative frequencies

Once the counts matrix is loaded (clusters on rows, samples on columns), the workflow is performed with the following code:

`# build the frequency matrix`

`freq_raw = sweep(counts, 2, colSums(counts), “/”)`

`# normalize: for each cluster, divide by its mean frequency freq_rel = sweep(freq_raw, 1, rowMeans(freq_raw), “/”)`

`# threshold relative frequency and convert to log2 thr = 4`

`freq_rel_lim = freq_rel # make a copy`

`freq_rel_lim[ freq_rel_lim > thr] = thr # replace high values freq_rel_lim[ freq_rel_lim < 1/thr] = 1/thr # replace low values log2_freq_rel_lim = log2(freq_rel_lim)`

`# heatmap`

`pheatmap(log2_freq_rel_lim, breaks = seq(−2, +2, length.out = 101))`

We performed a hard thresholding although we can specify color scale limits to pheatmap, which gets rid of the logarithm of zero problem. However, pheatmap needs finite data in order to automatically determine the data range to match the color scale or to calculate clustering.

Limiting the data range by hard thresholding removes the 0 from frequencies and also limits the impact of large differences on clustering. A few alternative solutions exist, but no consensus has been established to our knowledge (see Note 1).

In the differential analyses, one point is often overlooked: the reference group. When scaling frequencies to obtain relative frequencies, the average frequency of each cluster over all the samples is chosen. An interesting alternative is to select a group of samples in order to obtain a direct reading of changes with respect to this group and to evaluate the dispersion of this reference. Such a heatmap is presented on Figure 3C and could be compared with Figures 3A and 3B.

### 3.4 Building the cluster frequency heatmap and statistical analysis with Phantasus

#### 3.4.1 Importing counts

Open the Phantasus website, then select “My computer” in the “Choose a file” pop menu. Select the Excel file of counts in your computer. Phantasus presents the annotations and the numerical data matrix of the file. Rows annotations are highlighted in light green and are on the left of the sheet. Column annotations are highlighted in light red and are at the top of the sheet. The data are highlighted in light blue. Click the top left cell containing a count. The clusters must be on the rows and samples on the columns. Click “OK”.

#### 3.4.2 Performing the statistical analysis

Click “Tools / Differential expression / DESeq2” to perform the DESeq2 analysis via the menu. Enter “acute” in “Target level” and select all the 43 entries. Enter “conv” in “Reference level” and select all the 43 entries. When clicking “OK”, the results appear on the right for each cluster with log2FoldChange and padj (p-value adjusted for multi-testing) being the most important information for the interpretation. Don’t rely on the colors in the heatmap (Figure S1) as counts are normalized per row to the minimum and maximum.

#### 3.4.3 Computing the relative frequencies heatmap

To avoid zero counts, we add 1 to all the counts after calculating the total counts. Next, we calculate frequencies, their logarithms and subtract a reference mean, which is the same as calculating the relative frequencies and then their logarithm. See Note 5 to add a small value to frequency rather than counts. First, we calculate the total counts before adding 1. Select “Tools / Create Calculated Annotation”, tick “Columns”, click “Sum” in the “Operation” drop-down menu and “OK”. Then, we add 1 to the counts and calculate the frequencies. Select “Tools / Adjust”, tick “One plus log 2”, “Inverse log 2”, and in the “Sweep” section, select “Divide” each “column” by field “Sum”, and click “OK”. Next, we calculate the logarithm. Select “Tools / Adjust”, tick “Log 2” and click “OK”. Now, the values are log2 frequencies. To define the “Conv” sample group as a reference for relative frequencies, select all its columns and then select “Tools / Create Calculated Annotation”, tick “Rows”, click “Mean” in the “Operation” drop-down menu, tick “Use selected rows and columns only” and click “OK”. Then, select “Tools / Adjust”, and in the “Sweep” section, select “Subtract” from each “row” field “Mean”, and click “OK”. To tune the display to the effective scale of the data, we select “View / Options”, untick “Relative color scheme”. Verify that “Transform values” is set to “None” (see Note 8), set the “Minimum” to “-3” and the “Maximum” to “+3” and click “Close”.

#### 3.4.4 Clustering with Phantasus

To apply a clustering, select “Tools / Clustering / Hierarchical clustering”. In the popup window, select “Euclidean” as “Metric”, “Complete” as “Linkage method”, “Rows” as “Cluster” and click “Close” to apply. We can also sort the clusters by adjusted p-values by clicking on the padj column heading. The final heatmap is presented on Figure 4 (see Note 9 for exporting).

**Figure 4:**
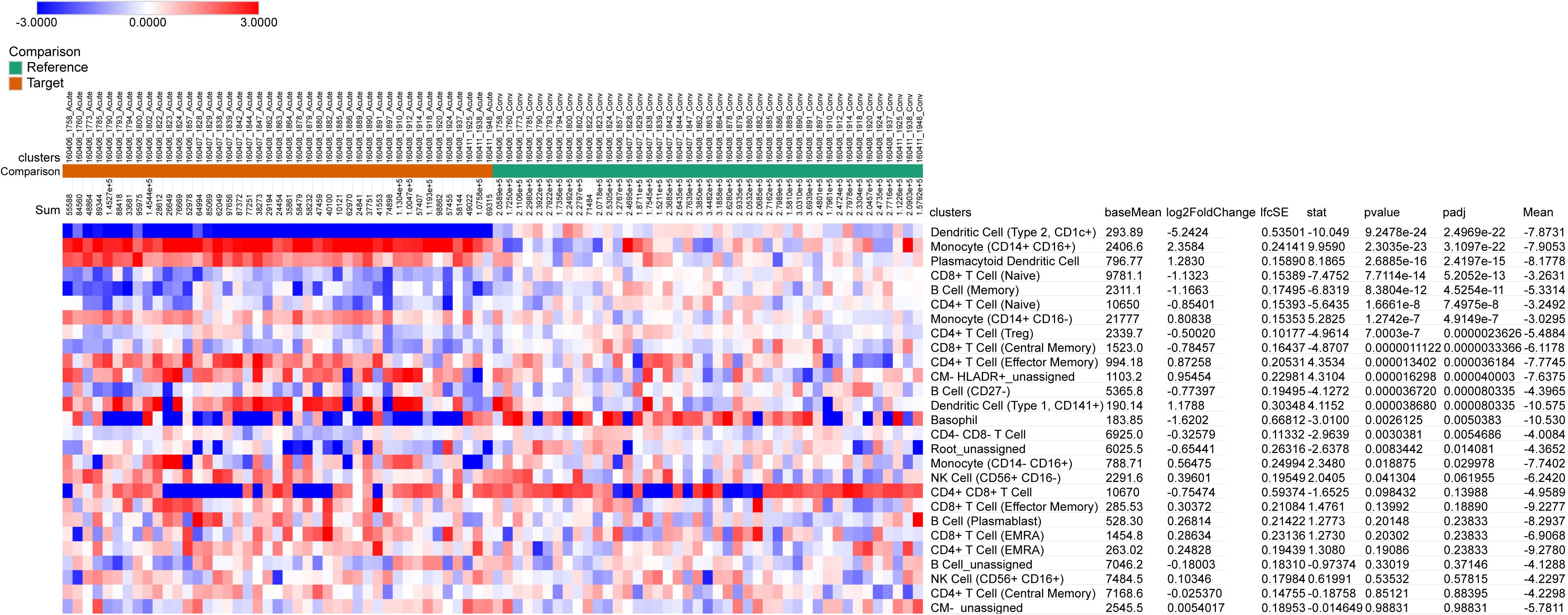
Final heatmap of frequencies relative to the “Conv” sample group along with the DESeq2 statistical analysis. As frequencies are relative to the “Conv” sample group, it is easier to interpret the changes and their strength by additionally relying on statistics.

## 4 Interpreting a heatmap

A heatmap is a visualization tool for exploring and interpreting data. To read a heat map and draw correct conclusions, we need to understand which object is in the row and which is in the column, to know how the color scale corresponds to the numbers, and to understand why and how the numbers have been processed.

The heatmap of transformed and scaled MFIs of markers per clusters aims to link clusters to cell populations. Markers are scaled independently to their minimum and maximum. The minimum must be defined (automatically or manually) with care, as its value misleads the reading of positively selected cells such as CD45+ cells. The maximum must also be carefully defined, as its value may overestimating the intensity of markers that are weakly expressed either natively or as a result of negative selection (as in figure 3 by Liechti at al. (11)). These two limits need to be particularly scrutinized by analyzing the colors in the heatmap for each marker and comparing them with our knowledge. Applying a clustering algorithm to rows and columns is crucial to facilitate the annotation of clusters, and its parameters should also be evaluated.

The heatmap of log2 relative frequencies of clusters across samples aims to identify or point out changes between groups of samples, which is directly related to the study question.

Changes are usually viewed and expressed as logarithm fold changes (logFC) and should be visualized with a symmetrical and limited color scale. When a group of samples is the reference in the experiment, it should be used to compute the relative frequencies instead of using all the groups. The heatmap then provides a direct reading of changes with respect to this group and of its dispersion. When the changes of relative frequencies are normalized to the standard deviation of the reference group (or the averaged within group deviations), the unit of the color scale is no longer the logFC, but a logFC relative to the dispersion, and must be interpreted cluster by cluster. So, even if the color scale is presented with numbers, knowing the heatmap building is important to correctly interpret its units. Applying a clustering algorithm to clusters makes it easier to group them together based similar changes over samples. Applying a clustering algorithm to samples makes it possible to verify the homogeneity of the groups studied or to detect subgroups.

We have presented the step-by-step construction of heatmaps used in cytometry. These workflows provide a clear understanding and enable the importance of steps and parameters to be assessed. However, R remains a more reproducible approach. A heatmap is not intended to perform statistical analyses but to complement them. Statistical analysis of counts (aka DA) requires a dedicated framework (5–7), in which raw counts are not transformed (Figure S1). A heatmap is a versatile visualization tool that covers many uses and can reveal or confirm interesting results of cytometry data, but understanding its building is mandatory.

## 5 Notes

1. We present a hard threshold of the relative frequency (to the chosen reference) before calculating the logarithm. An alternative well-known method is to add a prior of 1 to all the counts after calculating the total count per sample. Frequencies are then computed (with a denominator equal to the total count plus two times the prior in the empirical Bayesian framework). As long as the prior is small compared with the cell count the scientist considers convincing, and the total counts are much bigger, this method is a good approximation. Another method consists in adding a small value to the frequencies. In both methods, there is no rule for determining the additive value. Alternatively, all relative frequencies equal to zero can be replaced by the smallest non-zero value before applying the logarithmic transformation. There are probably other methods and transformations (square root, arcsine of the square root) that could be evaluated.
2. Phantasus can be used to carry out the whole workflow to obtain scaled transformed MFIs but with some cautions. Transforming raw MFI is usually performed using a bi-exponential or arcsinh function. The log2(x+1) transformation available in Phantasus may look similar for mass cytometry intensities that are typically lower than a few thousands, but we cannot control the width of the linear region before moving to the log region. This should be avoided, and the safe solution is to start with transformed intensities. To scale markers (columns), click on “Tools/Create Calculated Annotation”, then click “Columns” and select “Max” as “Operation”. Columns can then be normalized to their own maximum. Select “Tools/Adjust”, then at the bottom of the window in the “Sweep” section, select “Divide” each “Column” by field “Max”. Phantasus proposes a heatmap ready for interpretation, bearing in mind that no fine-tuning of markers minima and maxima could be achieved, which means that maxima may be too high.
3. Phantasus allows to perform statistical analyses. DESeq2 should be performed on the raw counts, without any adjustment, even if the heatmap appears suboptimal. See Fig S1.
4. When transforming values in log2 scale, Phantasus converts 0 into 0. As a result, any value below 1 becomes negative after transformation, even though it was originally greater than 0. This is a major problem when the values are frequencies (i.e. between 0 and 1 or between 0 and 100%). If there is no zero count, using Phantasus poses no problem for obtaining a correct heatmap. See 3.4 and Note 5.
5. Starting from the raw counts, we normalize these counts to a fixed total count for each sample and calculate the logarithm plus one. Select “Tools / Adjust”, tick “Scale column sum”, enter a target total count, here “30000” (because many samples have a low cell count of the order of tens of thousands and only 1 is added to this target count), tick “One plus log 2”, and click “OK”. Now, the values are log2 frequencies with a small value of 1/30000 added. Next, we can follow the workflow applied to the counts and define the reference to obtain log2 relative frequencies (Figure S2).
6. The shortcuts are valid for Excel under Windows. For Apple computer, the Ctrl key is replaced by the Apple key. As Excel translates the formula to your region, the words SUM, AVERAGE, IF must be replaced the corresponding word in your language. The list separator in formulas should also be replaced by “,” instead of “;”.
7. Excel files are interesting because the numbers are encoded in a format that does not depend on location. The decimal separator is displayed when numbers are displayed and adapted to localization, but their internal encoding is unique and portable. The same applies to date formats, but it is advisable to use the YYYY-MM-DD format, which is an unambiguous international standard.
8. Phantasus (and other software) transforms the data on-the-fly to derive a value that lies within the same range for the entire dataset. This leads to two datasets, the imported dataset itself (and possibly processed using “Tools/Adjust”) and the dataset resulting from the on-the-fly transformation for coloring. Nevertheless, Phantasus still presents the actual data in the tooltip when the mouse is moved over the heatmap. This duality can be misleading and we prefer to avoid it in order to visualize the result of a transformation on the data. Phantasus offers on-the-fly transformations that apply to rows (typical of Omics analyses), which implies that the dataset is adequately prepared (transposition is available during the import). By default, the “Relative color scheme” is selected, using the minimum and maximum of each row to adjust the color scale per row. By unticking this option, two transformations are possible. “Subtract row mean, divide by row standard deviation” is the classical statistical standardization. “Subtract row median, divide by row median absolute deviation” is the alternative standardization robust to outliers. These two transformations could be interesting when we first discover the dataset, but scaling and color scaling are then specific to each row.
9. To export a nice figure from Phantasus, click on the icon showing a landscape (or use the menu “File / Save image”), then select either PNG for a screenshot or SVG (or PDF) for a scalable and editable graph. The exported figure will comprise the color scale and the legend of annotations. The color scale takes into account an on-the-fly transformation. If the default row min max scaling is activated, the limits of the color scale are “row min” and “row max” instead of numerical values when transformation is “None”. The figure can be improved by removing the grid and adjusting the column width.
10. To load the prepared R data files, use the following commands: `load(”mfis_Assignment.RData”) load(”counts_Assignment.RData”)`

To import the counts matrix from an Excel file, use the following code that requires the installation of the readxl package beforehand. Use a similar code for MFIs.

`tmp = as.data.frame(readxl::read_excel(”counts_Assignment_matrix.xlsx”))`

`counts = as.matrix(tmp[,-1]) rownames(counts) = tmp[,1]`

---

**Figure S1:**
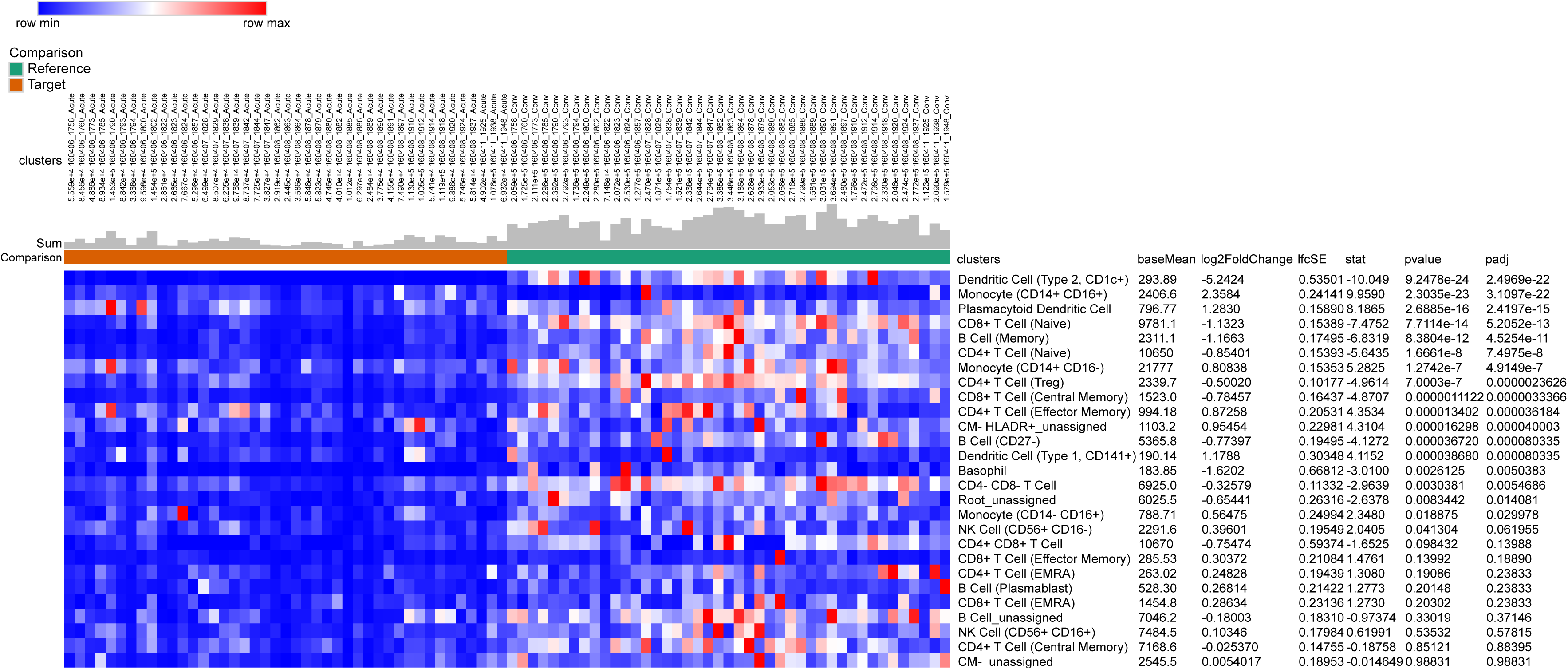
DESeq2 analysis with Phantasus. Counts were imported and directly subjected to a DESeq2 analysis via the menu “Tools / Differential expression / DESeq2”. Clusters (identified cell populations) are sorted by increasing adjusted p-value. The colors in the heatmap are normalised per cluster (row) to the minimum and maximum by default. The coloring is biased by the fact that there are more cells in the “Conv” samples than in the “Acute” ones. The statistical analysis clearly takes this bias into account, as “Monocyte (CD14+ CD16+)” cluster is statistically significant but not detectable in the heatmap. The sum cells per sample (column) was added using “Tools / Created Calculated annotation” and displayed with 4 digits and as a bar chart.

**Figure S2:**
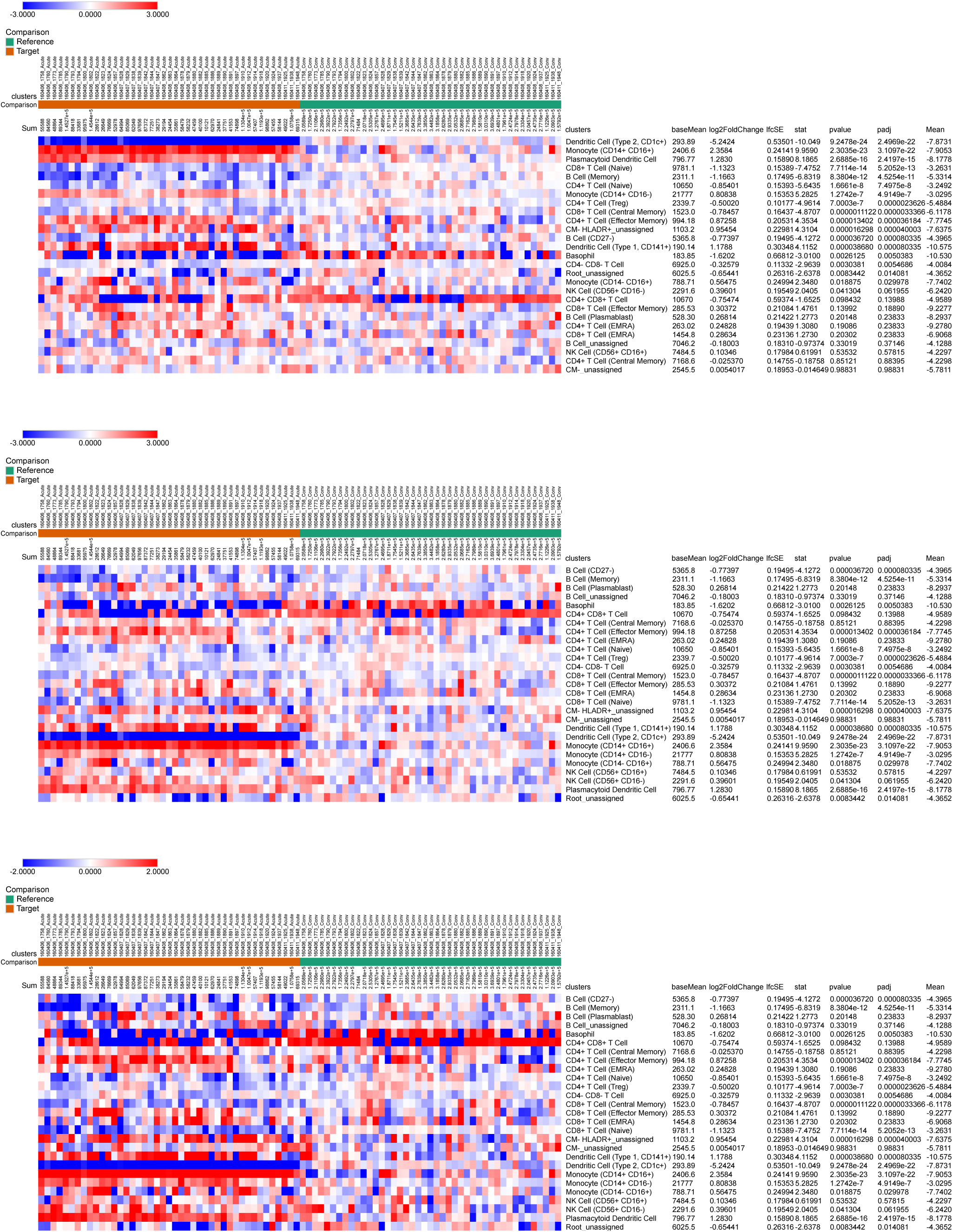
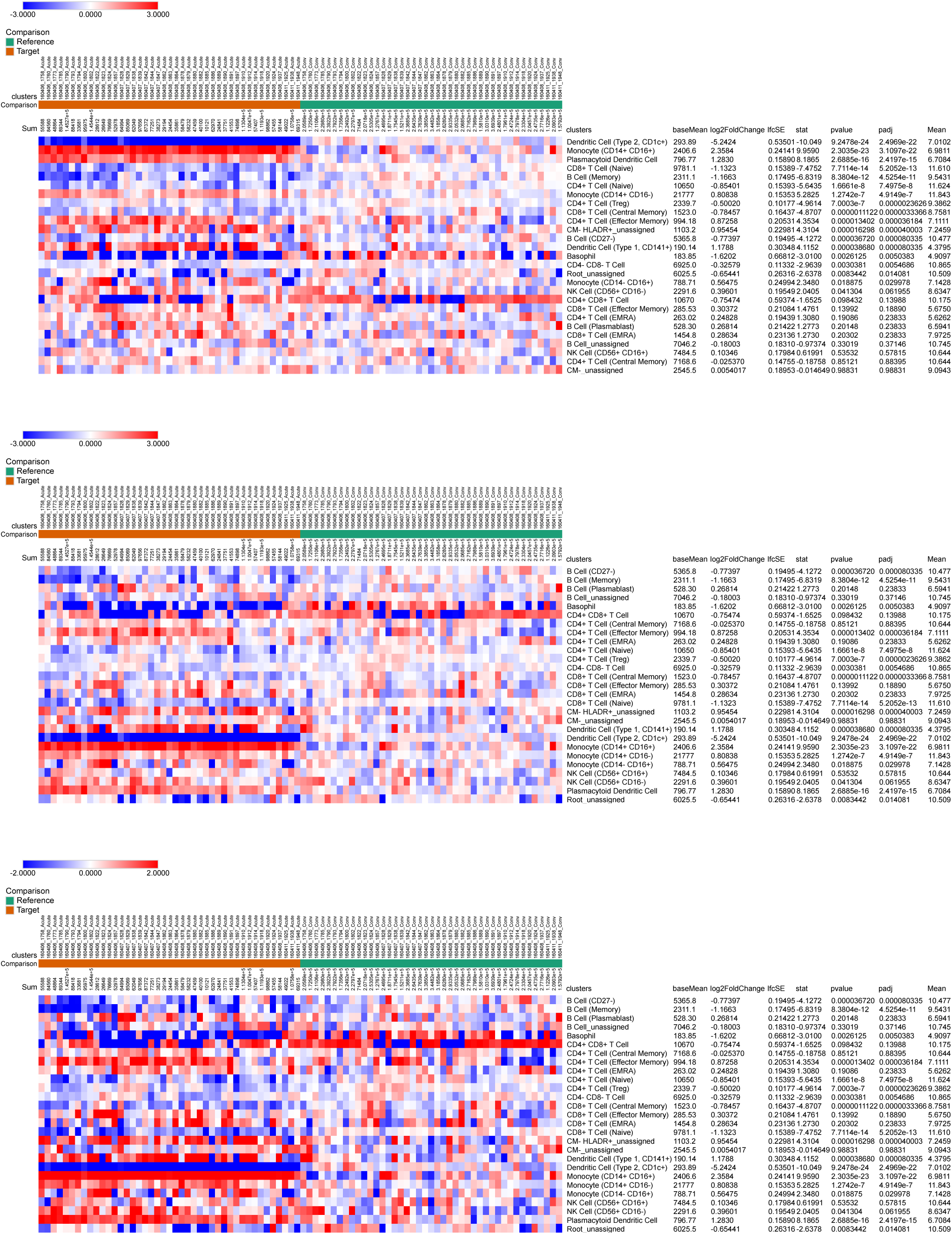
Comparison of count processing: count + 1 workflow versus freq + 1/30000 workflow.

## References

1. Eisen MB, Spellman PT, Brown PO, Botstein D (1998) Cluster analysis and display of genome-wide expression patterns. Proc Natl Acad Sci U S A 95:14863–14868. 10.1073/pnas.95.25.14863

2. Saeed AI, Sharov V, White J, et al (2003) TM4: a free, open-source system for microarray data management and analysis. Biotechniques 34:374–378. 10.2144/03342mt01

3. Kleverov M, Zenkova D, Kamenev V, et al (2024) Phantasus, a web application for visual and interactive gene expression analysis. eLife 13:e85722. 10.7554/eLife.85722

4. Weber LM, Nowicka M, Soneson C, Robinson MD (2019) diffcyt: Differential discovery in high-dimensional cytometry via high-resolution clustering. Commun Biol 2:1–1. 10.1038/s42003-019-0415-5

5. Robinson MD, McCarthy DJ, Smyth GK (2010) edgeR: a Bioconductor package for differential expression analysis of digital gene expression data. Bioinformatics 26:139–140. 10.1093/bioinformatics/btp616

6. Chen Y, Chen L, Lun ATL, et al (2025) edgeR v4: powerful differential analysis of sequencing data with expanded functionality and improved support for small counts and larger datasets. Nucleic Acids Res 53:gkaf018. 10.1093/nar/gkaf018

7. Love MI, Huber W, Anders S (2014) Moderated estimation of fold change and dispersion for RNA-seq data with DESeq2. Genome Biol 15:550. 10.1186/s13059-014-0550-8

8. Diggins KE, Gandelman JS, Roe CE, Irish JM (2018) Generating Quantitative Cell Identity Labels with Marker Enrichment Modeling (MEM). Curr Protoc Cytom 83:10.21.1-10.21.28. 10.1002/cpcy.34

9. Kolde R (2019) pheatmap: Pretty Heatmaps https://cran.r-project.org/package=pheatmap

10. Michlmayr D, Pak TR, Rahman AH, et al (2018) Comprehensive innate immune profiling of chikungunya virus infection in pediatric cases. Mol Syst Biol 14:e7862. 10.15252/msb.20177862

11. Liechti T, Weber LM, Ashhurst TM, et al (2021) An updated guide for the perplexed: cytometry in the high-dimensional era. Nat Immunol 22:1190–1197. 10.1038/s41590-021-01006-z

